# HDAC inhibition protects degenerating cone photoreceptors *in vivo*

**DOI:** 10.1101/049742

**Authors:** Dragana Trifunović, Blanca Arango-Gonzalez, Antonella Comitato, Melanie Barth, Ayse Sahaboglu, Eva M. del Amo, Manoj Kulkarni, Stefanie M. Hauck, Marius Ueffing, Arto Urtti, Yvan Arsenijevic, Valeria Marigo, François Paquet-Durand

## Abstract

Retinal diseases caused by cone photoreceptor cell death are devastating as the patients are experiencing loss of accurate and color vision. Understanding the mechanisms of cone cell death and the identification of key players therein could provide new treatment options. We studied the neuroprotective effects of a histone deacetylase inhibitor, Trichostatin A (TSA), in a mouse model of inherited, primary cone degeneration (*cpfl1*). We show that HDAC inhibition protects cones *in vitro*, in retinal explant cultures. More importantly, *in vivo* a single TSA injection increased cone survival for up to 10 days post-injection. In addition, the abnormal, incomplete cone migration pattern in the *cpfl1* retina was significantly improved by HDAC inhibition. These findings suggest a crucial role for HDAC activity in primary cone degeneration and highlight a new avenue for future therapy developments for cone dystrophies and diseases associated with impaired cone migration.

## Introduction

Cone photoreceptors in the human retina are responsible for sharp high resolution vision and color discrimination. Hereditary cone degenerations, such as in Stargardt and Best disease, achromatopsia, and cone dystrophies are caused by mutations in single genes, and lead to severe visual impairment, reduced visual acuity, and loss of color vision. Cone dystrophies are characterized by a high genetic heterogeneity with a large variety of mutations in at least 27 genes (RetNet: https://sph.uth.edu/retnet). In addition, genetic causes have also been proposed for complex diseases affecting cones, such as in age-related-macular degeneration (AMD) or diabetic retinopathy (DR). Irrespective of the genetic causes, a common outcome of retinal diseases is neuronal cell death, giving a strong rationale for targeted neuroprotective approaches that may prevent or delay cell death execution ^1^.

The c*one photoreceptor function loss-1* (*cpfl1*) mutant mouse is an animal model for autosomal recessive achromatopsia or progressive cone dystrophy ^2^. The *cpfl1* mouse model is unique in the sense that it is characterized by an early onset of cone loss at post-natal day 14 (PN14) and a fast progression with the peak of cell death at PN24 ^3^. We have previously shown that photoreceptor cell death in the *cpfl1* mouse, as well as in nine other animal models for inherited retinal degeneration, follows a non-apoptotic mechanism, characterized by accumulation of cGMP, increased activities of cGMP-dependent protein kinase (PKG), histone deacetylase (HDAC), poly-ADP-ribose-polymerase (PARP), and calpain proteases, as well as accumulation of poly-ADP-ribose (PAR) ^4^. In addition, the *cpfl1* cone degeneration is also associated with prominent defects in cone migration ^3^.

A common non-apoptotic cell death mechanism in different retinal degeneration models provides a rationale for the identification of therapeutic targets that would prove beneficial for a broader population of patients suffering from a variety of different genetic causes ^4^, ^5^. While, numerous attempts to restrain the execution of rod photoreceptor cell death were described ^5^, ^6^, ^7^, ^8^, ^9^ to date, an effective clinical treatment for primary cone photoreceptor degeneration has not been reported. Since humans use mostly cones for their vision, a development of treatment protocols to preserve cones is of highest priority in clinical ophthalmology.

The interplay between histone acetylation and deacetylation, performed by histone acetyltransferases (HATs) and histone deacetylases (HDACs) respectively, determines the transcription state of genes in general^10^, and this is also true for photoreceptor-specific genes ^11^, ^12^. Aberrant activity of histone deacetylases is associated with a number of diseases with very different etiology, ranging from cancer to neurodegenerative diseases ^13^. Consequently, in current medicine, HDAC inhibition is discussed as one of the most promising novel therapeutic approaches for various diseases ^14^.

In the present study, we tested the hypothesis that hereditary cone degeneration can be prevented or delayed by pharmacologically inhibiting HDAC. We assessed cone photoreceptor survival after HDAC inhibition with Trichostatin A (TSA) *in vitro* on retinal explant cultures obtained from *cpfl1* animals, as well as *in vivo* after intravitreal injection. We show that a single TSA injection *in vivo* achieved protection of cone photoreceptors, up to ten days post-injection. In addition, we observed a significant improvement of impaired cone migration, present in degenerating *cpfl1* retina. Our study shows for the first time the possibility to use pharmacological HDAC inhibition for massive cone protection in an inherited cone dystrophy. These results may be highly relevant for the future development of therapies aimed at cone photoreceptor preservation.

## Results

### HDAC activity is increased in *cpfl1* photoreceptors

We have previously shown that the *cpfl1* cone photoreceptor degeneration follows a non-apoptotic cell death mechanism characterized by increased HDAC activity at the peak of degeneration at PN24 ^4^. In this study, we evaluated HDAC activity also at the onset of cone degeneration (PN14). As soon as *cpfl1* cone degeneration was detectable by TUNEL staining, we found significantly more HDAC activity positive photoreceptors in mutant retinas compared to wild-type (wt) (see Supplementary Fig. S1). The percentage of cells positive for HDAC activity in the photoreceptor layer corresponded to the percentage of dying photoreceptors at PN14, suggesting that increased HDAC activity was related to cone degeneration.

### HDAC inhibition with TSA rescues degenerating cones *in vitro*

To test whether HDAC inhibition could promote cone photoreceptor survival, we used *cpfl1* long-term organotypic retinal explant cultures as *in vitro* system. Both *cpfl1* and wt retinas were explanted at the onset of *cpfl1* degeneration (PN14) and cultured for 2 days without any treatment. Explanted retinas were then exposed to 10 nM TSA treatment every second day, until PN24. TSA had no effect on cone photoreceptors in wt cultures, as assessed by the percentage of cones in the outer nuclear layer (ONL), identified by labeling with an antibody for glycogen phosphorylase (Glyphos) (TSA: 4.62% ± 1.98 SEM, n=3; untreated: 4.2% ± 0.55, n=4; Figure 1). In the *cpfl1* retina, however, 10 nM TSA significantly increased the percentage of surviving cones from 2.21% ± 0.32 SEM (n=7) to 4.6% ± 0.47 (n=6, *p= 0.0013*; Figure 1), which corresponded to an increase from 47.6% to 99.5% cones, when compared to the wt situation. Hence, HDAC inhibition appeared to afford a full protection of *cpfl1* cones.

**Figure 1.**
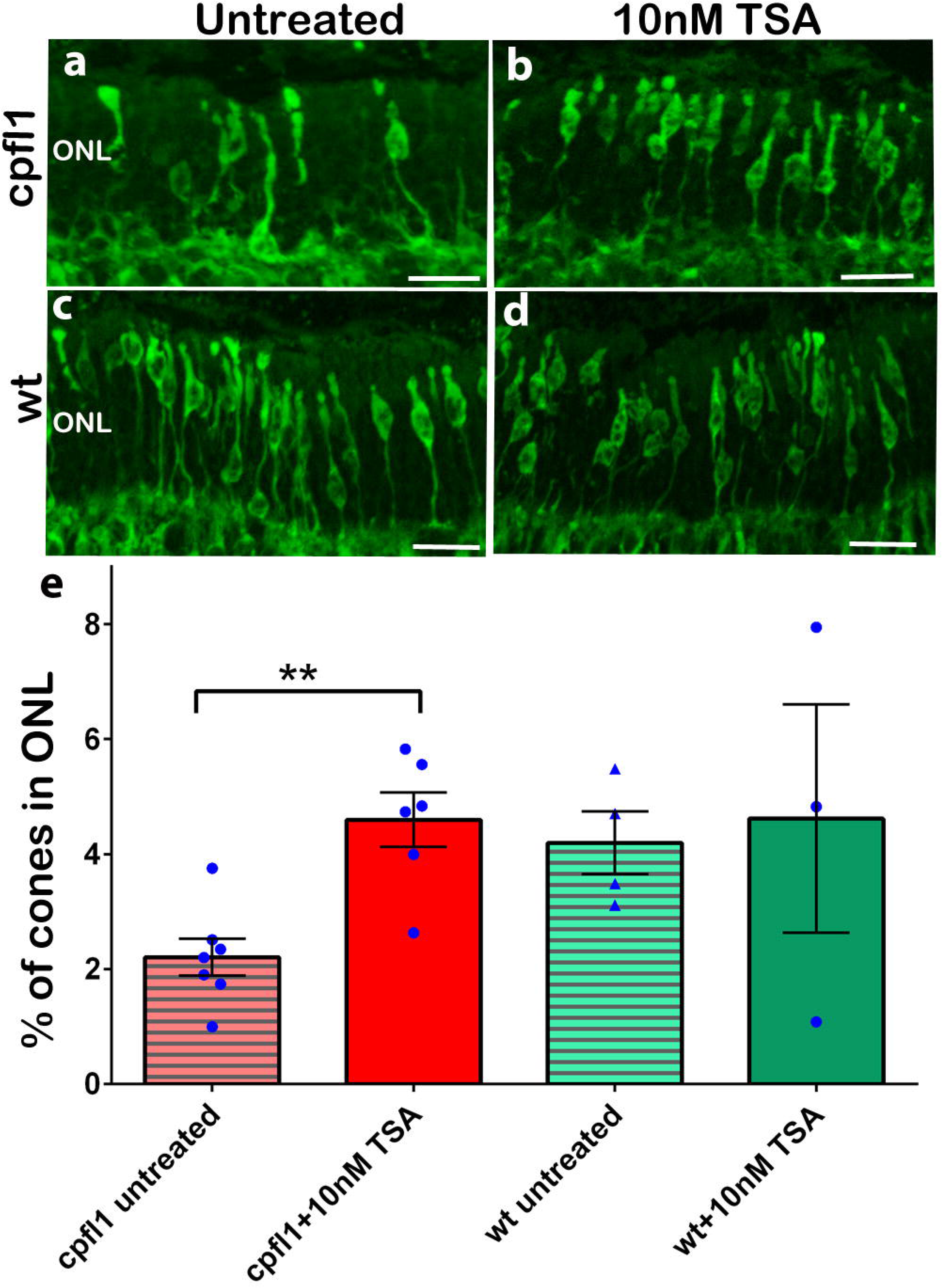
HDAC inhibition protects cone photoreceptors *in vitro*. Long term retinal explant cultures from *cpfl1* and wt animals were subjected to 10 nM TSA or control condition from PN14 to PN24. Glyphos staining (green), specific for cone photoreceptors, suggested that very few cones survived at PN24 in untreated *cpfl1* explant cultures (a), while TSA treated retinas displayed prominently increased numbers of cones (b). TSA treatment showed no effect on wt treated retinas (c, d). The protective effect of TSA on cones is summarized in the bar graph (e). Error bars represent SEM; Scale bars are 20 μm. ONL-outer nuclear layer containing photoreceptors. INL-inner nuclear layer containing interneurons. GCL-ganglion cell layer.

### Intravitreal TSA injection protects degenerating cones *in vivo*

The promising *in vitro* results prompted us to also engage in a study to evaluate the protective effects of HDAC inhibition *in vivo*. To optimize the *in vivo* treatment scheme, we first tried to predict the potential half-life of the drug inside the eye. Model calculations suggested that intravitreal clearance of TSA in the rabbit eye is 0.478 ml/h, and the half-life is in the range of 1.7-3.3 hours. In the mouse, the calculated intravitreal half-life of TSA is an order of magnitude shorter (about 17 minutes), and more than 90% of the dose is expected to be eliminated in one hour after injection.

Since our *in vitro* experiments suggested that TSA was efficiently protecting cones in concentrations as low as 10 nM, we decided to test if the same concentration would be effective also *in vivo*. As controls, we included both non-treated and sham treated eyes in the analysis. At PN14, nine animals from three different litters received an injection of 0.5 μl of a 100 nM TSA solution in one eye, with an estimated final TSA concentration of 10 nM. The contralateral eyes were sham injected to control for injection-specific effects. Mice were sacrificed and analyzed after 10 days, at PN24. Consistent with our *in vitro* results, TSA significantly increased the percentage of surviving cones in treated *vs*. sham treated eyes (*cpfl1* sham: 4.57% ± 0.18 SEM, n=9; *cpfl1* treated: 5.37% ± 0.27, n=9, *p= 0.027*; Fig. 2). The percentage of cones in sham injected eyes was comparable to the percentage of cones in non-treated *cpfl1* mice (*cpfl1*: 4.03% ± 0.23 SEM, n=9; *cpfl1* sham: 4.57% ± 0.18, n=9), suggesting that the injection itself had no effect on cone survival. The number of cones in sham- or 10 nM TSA- treated wt retinas were similar to non-treated wt retinas confirming the *in vitro* observation that TSA in the studied dosage had no deleterious effect on healthy cones (wt: 5.38% ± 0.38 SEM, n=5; wt sham: 5.43% ± 0.26, n=8; wt treated: 5.71% ± 0.38, n=8, Fig. 2). The *in vivo* results suggested a remarkable protection of degenerating cones by a single intravitreal injection of 10 nM TSA with the number of cones in treated retinas reaching wt levels, *i.e. cpfl1* treated: 5.374% ± 0.78 *vs*. wt: 5.378% ± 0.38.

**Figure 2.**
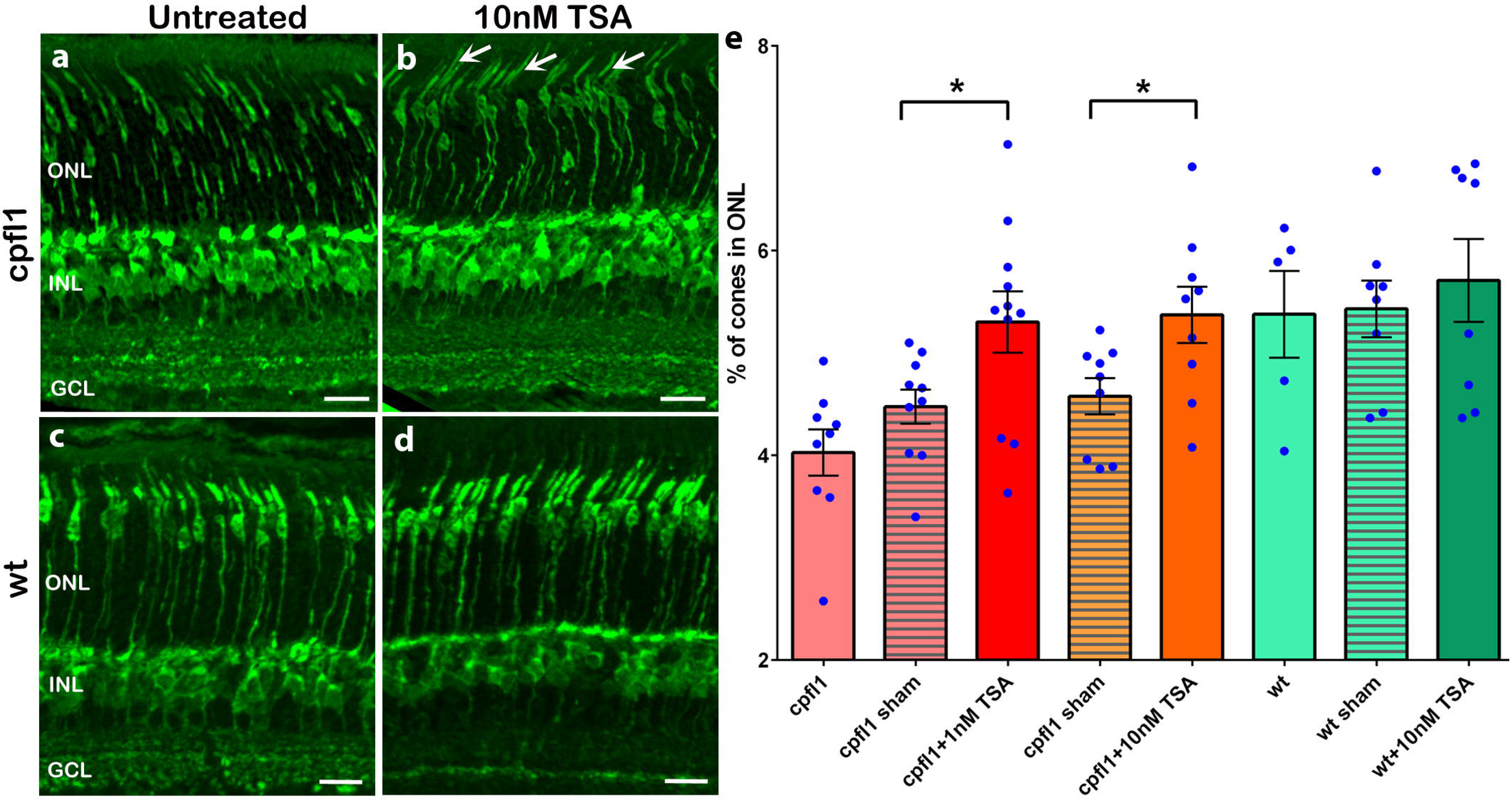
HDAC inhibition protects degenerating cones *in vivo*. A single intravitreal injection of 1 nM or 10 nM TSA at PN14 was sufficient to significantly increase the number of *cpfl1* mutant cones in treated compared to sham injected retinas at PN24 (a, b). In addition, cones from treated retinas were positioned mainly in the upper part of the ONL and had visibly longer inner segments (arrows) in comparison with sham treated eye. *In vivo* treatment of wt animals showed no effects on the cone number, cone positioning, or structure (c, d). Quantification of the cone percentage of non-treated, sham treated and TSA treated (1 nM and 10 nM) *cpfl1* and wt animals is shown in the bar graph (e). Remarkably, the percentage of cones present was similar to wt for both TSA concentrations. Scale bar in a-d is 20 μm.

We then tested if TSA would efficiently protect degenerating cones at lower concentrations. We performed a single intravitreal injection of estimated 1 nM TSA at PN14 and evaluated the protective effects at PN24. Remarkably, 1 nM TSA single treatment was sufficient to fully protect cones in *cpfl1* retina (Fig. 2, *cpfl1* sham: 4.48% ± 0.16 SEM, n=10; *cpfl1* treated: 5.3% ± 0.29, n=11, *p=0.03*). Wild-type retinas treated with 1 nM TSA showed no changes in the number of cones compared to sham treated (wt sham: 5.58% ± 0.43, n=3; wt treated: 5.39% ± 0.4, n=3). Since the protective effect of TSA might have been due to a delay of cone maturation in *cpfl1* animals it was important to characterize the status of cone differentiation. To this end, we looked for the expression and localization of M- and S-opsins as well as cone-specific transducin (GNAT2), previously reported to be abnormally distributed in degenerating cones ^15^. Although M-opsin is present in *cpfl1* cone outer segments, several cones showed opsin mislocalization in the cell body (Fig. 3g, arrows) and photoreceptor end feet (Fig. 3g, arrowheads). A similar but more pronounced mislocalization was observed for GNAT2 (Fig. 3i, arrows) in *cpfl1* retina. S-opsin staining revealed short outer segments as well as opsin mislocalization (Fig. 3h, arrows). Double staining for M- and S-opsin with Glyphos indicated that TSA protected cones had normal morphology, with clearly developed inner and outer segments (Fig. 3p-r). In addition, the correctly positioned cones in the upper part of the ONL expressed both M- and S-opsin in their outer segments. Similarly, in surviving cones GNAT2 was properly localized in outer segments. The correct localization of cone opsins and cone transducin suggested that both 1nM and 10nM TSA had preserved cones, without adverse effects on cone development and maturation. Thus, further research on different TSA dosage effects on cone survival is warranted.

**Figure 3.**
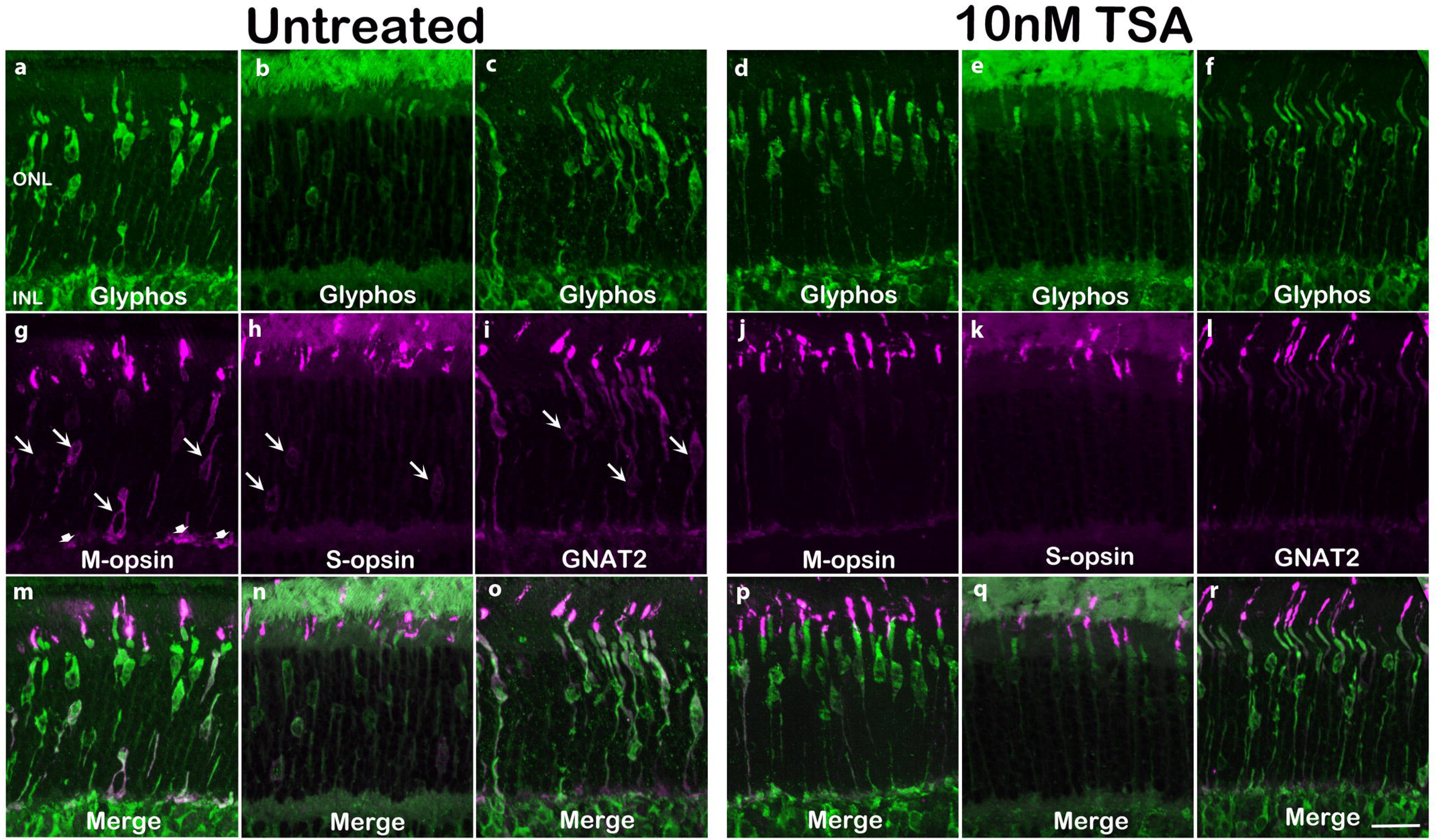
Cone specific opsins and transducin are expressed and positioned correctly in protected cones. In untreated PN24 *cpfl1* retinas, M-opsin (red color, g, m) was correctly localized in outer segments in some cones stained with Glyphos (green, a, m), mainly in those positioned in the upper parts of the ONL. At the same time, M-opsin was also mislocalized in what seemed to be cytoplasm of misplaced cones close to the border with the INL (arrows) and in cone end feet (arrowheads). A similar mislocalization was observed also with antibodies directed against S-opsin (b, h, n) and transducin (GNAT2; arrows; c, i, o). However, after TSA treatment, not only were there more cones, but M-opsin, S-opsin, as well as GNAT2 were properly localized almost exclusively in outer segments (p-r). Scale bar is 20 μm.

### HDAC inhibition significantly improves impaired cone migration

We previously reported that the *cpfl1* retina is characterized by impaired migration of developing cones ^3^. Glyphos staining of cones in degenerating *cpfl1* retina confirmed such aberrant cone positioning, both *in vitro* and *in vivo* (Fig. 4). Importantly, we observed an improved migration of cones after both *in vitro* and *in vivo* TSA treatments of retinas (Fig. 4). To evaluate the extent of the cone migration improvement, we measured the distance of the center of individual cone nuclei from the outer plexiform layer (Fig. 4, dashed line), which demarcates the lower boundary of the photoreceptor area. The percentage of cone migration distance from the OPL is presented in relative terms to the thickness of the ONL. In the *in vitro* situation, at PN24, wt cones migrated approximately to 70% of ONL thickness (wt untreated: 69.25% ± 2.3 SEM, n=3; wt treated: 72.66% ± 5.1, n=3), while in non-treated *cpfl1* retinas they reached only 55%. After treatment with 10 nM TSA of *cpfl1* retinas in explant cultures, cone nuclei reached a position very similar to that of wt cones (*cpfl1* untreated: 55.31% ± 3.1 SEM, n=6; *cpfl1* treated: 65.95% ± 2.8, *p=0.029*). Similarly, cone migration was also considerably improved in *cpfl1* retinas *in vivo*, after treatment with 1 nM TSA. Here, cones were localized significantly more distant from the outer plexiform layer (*cpfl1* untreated: 71.68% ± 1.09 SEM, n=10, *cpfl1* treated: 77.18% ± 0.84, n=11, *p=0.02*). Comparable results were obtained after treatment with 10 nM TSA (*cpfl1* untreated: 67.47% ± 1.92 SEM, n=9, *cpfl1* treated: 76.38% ± 0.73, n=9, *p=0.0005*). We did not observe changes in cone localization in wt retinas after TSA treatment (wt untreated: 86.89% ± 0.26 SEM, n=7; wt treated: 85.49% ± 0.23, n=7). Nevertheless, even though TSA treatment significantly improved *cpfl1* cone migration, cone positioning was still significantly different compared to wt cones (*cpfl1* 1nM TSA: 77.18% ± 0.84 SEM *vs*. wt untreated: 86.89% ± 0.26, *p=0.0002*; *cpfl1* 10nM: 76.38% ± 0.73 *vs*. wt untreated: 86.89% ± 0.26, *p=0.0004*).

**Figure 4.**
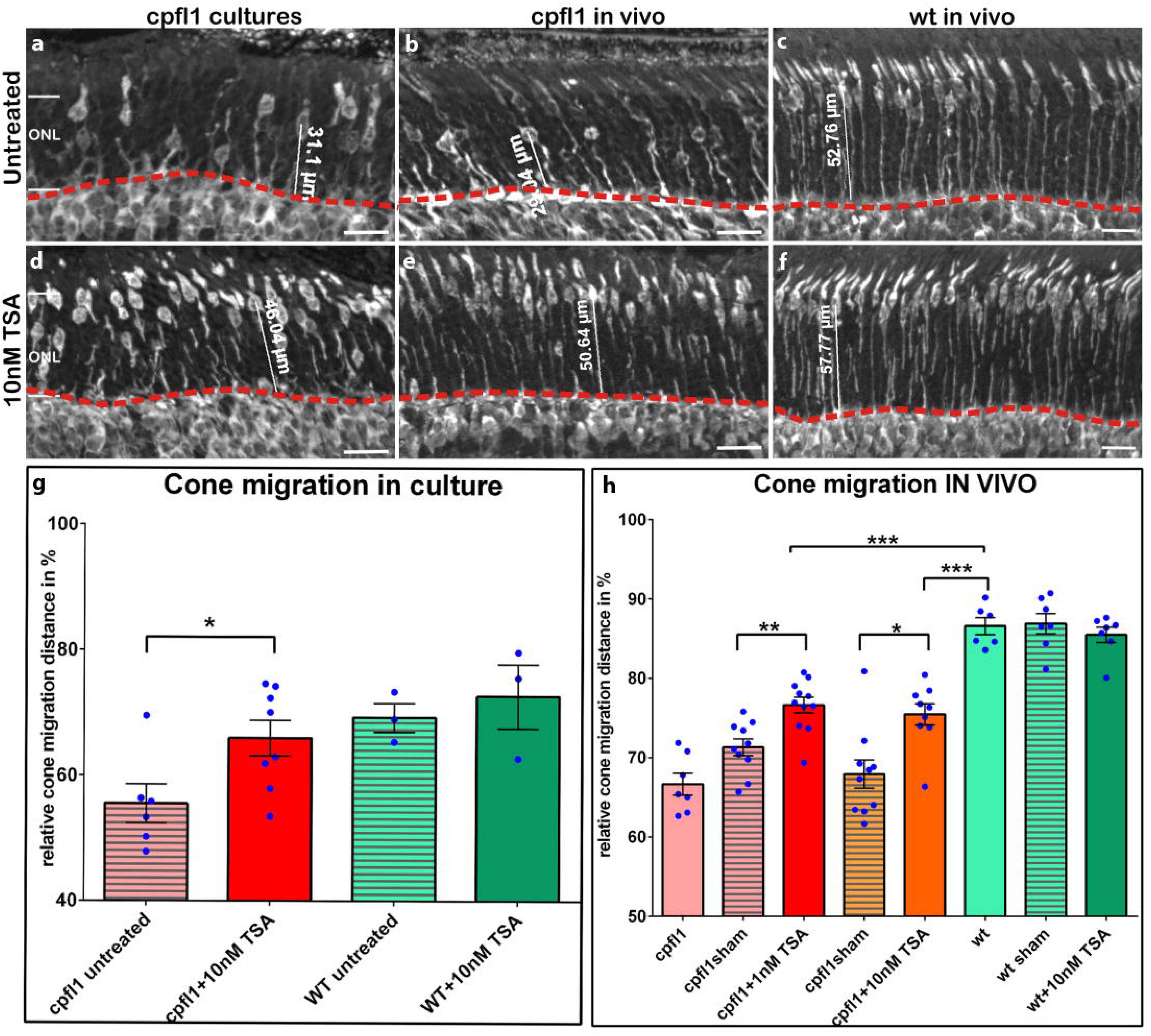
HDAC inhibition improves cone migration. Cone staining in *cpfl1* retina revealed a cone mislocalization, when compared to wt (a-c). Cone nuclei were scattered throughout the ONL in *cpfl1* mutant retinas also after culturing (a), while in wild-type retinas cones are positioned exclusively in the upper part of the ONL (c). TSA treatment significantly improved cone migration, both in the *in vitro* (d) and *in vivo* (e) treatment paradigms, while no effect was detectable in wt treated retinas (f). Cone migration distance was assessed by measuring the distance between the center of cone nuclei from the outer plexiform layer (dashed line), relative to ONL thickness, measured for each individual section of analyzed retinas. Scale bars are 20 μm.

We have previously found high cGMP accumulation and activity of cGMP-dependent PKG to be associated with *cpfl1* cone death as well as with cone mislocalization ^3^. To investigate whether HDAC inhibition at early PN14 had an effect on cGMP accumulation and PKG activity after ten days of treatment, we looked for cGMP accumulation and the phosphorylation status of a well-known PKG substrate, vasodilator-stimulated-protein (pVASP), with and without TSA treatment. Untreated *cpfl1* retinas showed cGMP accumulation in cone segments as well as in cone cell bodies (as evidenced by Glyphos co-labeling). cGMP accumulation could still be detected in cone segments after TSA treatment but was not observed in cone cell bodies (see Supplementary Fig. S3a). Likewise, some cones, visualized by Glyphos staining, were also positive for pVASP in both TSA treated and untreated *cpfl1* retinas, with no evident differences between the two (Fig. S3b).

## Discussion

Hereditary diseases of cone photoreceptors lead to major visual impairment, blindness, and are currently untreatable. Our study provides new evidence that increased HDAC activity is causally involved in hereditary cone photoreceptor degeneration. Consequently, HDAC inhibition resulted in long-lasting cone protection *in vitro* and *in vivo*.

HDAC inhibition has been demonstrated to induce cell death in tumor cells derived from various cancers ^16^, while in neurodegenerative diseases, the same treatments may prolong cell survival in postmitotic neurons ^17^. In the retina, HDAC inhibitors have been discussed as a potential therapeutic strategy for ischemic retinal degeneration, where HDAC inhibition with different inhibitors afforded structural and functional neuroprotection in a model of ischemic retinal injury ^18^. Furthermore, HDAC inhibition was shown to slow rod photoreceptor degeneration in an animal model for Retinitis Pigmentosa ^19^. In *cpfl1* retina, a previous study found an upregulation of STAT3 signaling which was suggested as an endogenous neuroprotective response ^20^. Interestingly, STAT3 activation may be controlled by HDAC activity ^21^. We show here for the first time that HDAC inhibition almost fully protects cones in a hereditary retinal dystrophy. Besides its implication for future treatment of rare forms of retinal degeneration, the results presented here may also extend to common diseases of the retina, including diabetic retinopathy and age related macular degeneration, where cone degeneration is the cause of legal blindness. In addition, even though gene therapy is currently the only approach to significantly improve vision in patients suffering from hereditary retinal degenerations, photoreceptor cell death in treated patients is not prevented ^22^. This implies a need for a combined therapy to repair both the genetic defect and to preserve neuronal viability. In *cpfl1* mice, a previous study showed that interference with thyroid hormone signalling preserved cones to some extent ^23^. Significantly, this protective effect may have been dependent on the inhibition of thyroid hormone T3 binding to HDAC, which can also be disrupted by TSA ^24^. However, TSA affects HDAC more directly than anti-thyroid treatment, which may explain the fact that we see almost full cone protection, for the first time in a model for hereditary cone degeneration. Importantly, TSA did not need repeated administration and a single *in vivo* injection was sufficient to halt degeneration with a long lasting effect, as the protection was present even ten days post-injection. This is unexpected, as our calculations on TSA clearance suggested that the drug was below threshold concentration, after less than 2 hours and point to an imprinting mechanism, beyond the transient presence of TSA. This is in line with previous studies which found that TSA leads to robust protein acetylation after 3 hours ^11^ or to transcriptional changes after only 5 minutes ^25^, supporting the possibility of epigenetic mechanisms. Importantly, TSA clearance estimation suggested that cone degeneration was halted with a single, short-time intraocular exposure. This observation could be relevant in light of minimizing potential toxic effects observed in some cases of long-term, systemic HDAC inhibition ^26^, ^27^. Cone photoreceptor development is characterized by a migration within the ONL, from the outer limiting membrane inwards to the OPL and then outwards until they reach their final position, just below the outer limiting membrane, from PN12 onwards ^28^. It has previously been reported that cone degeneration in mouse models is characterized not only by cone loss, but also by improper developmental cone migration ^3^, ^29^. In fact, delayed cone migration was also reported in mouse models with rod degeneration as an indirect consequence of rod cell death ^28^. Migration of cone photoreceptors is also present in human retinal development, where cones are migrating laterally towards the foveal pit ^30^. Cone misplacement was also associated with human retinal diseases of different etiology as improper cone migration can lead to foveal hypoplasia ^31^, an abnormal foveal structure. Abnormal foveal structure is observed in patients suffering from albinism ^32^, idiopathic congenital nystagmus ^31^, and also from achromatopsia ^33^. In addition, cone misplacement was also reported in patients suffering from age-related macular degeneration ^34^. We found that TSA treatment not only prevented cone degeneration but also significantly improved aberrant cone migration in degenerating *cpfl1* retina. There are numerous reports on the effects of HDAC inhibition on cell migration. The effects of HDAC inhibition on improved cell migration could take place via inhibition of the cytoplasmic histone deacetylase (HDAC6) resulting in higher acetylation of α-tubulin necessary for stability of microtubules in migrating cells ^35^. Alternatively, HDAC inhibition could lead to epigenetic changes governing migration of photoreceptors, a process still not fully understood. TSA treatment significantly improved the migration of misplaced *cpfl1* cones both *in vitro* and *in vivo*. Nevertheless, HDAC inhibition alone was not sufficient to fully repair the cone migration defect in *cpfl1* retinas. A plausible explanation could relate to increased PKG activity in *cpfl1* retina, as we previously reported ^3^. PKG-specific phosphorylation of the serine 239 residue of VASP was shown to negatively regulate neuronal migration ^36^. Proteins of the Ena/VASP family are regulators of actin assembly and cell motility as they are localized in focal adhesions, dynamic membrane structures important for cell migration ^37^. cGMP-dependent PKG overactivation leads to VASP phosphorylation, which will result in removal of VASP from focal adhesions, contributing to altered cell migration ^38^. Our data suggest that TSA inhibition of HDACs did not reduce VASP phosphorylation of serine 239 as assessed by immunostaining. The possible dual regulation of cone photoreceptor migration, one via PKG overactivation and the other involving aberrant HDAC activity, will require further specific studies.

Degenerating cones in *cpfl1* retina are characterized by cGMP accumulation as a consequence of non-functional *Pde6c* ^3^, ^4^. We did not observe obvious changes in the numbers of cells showing cGMP accumulation after the treatment, indicating that TSA-driven protection of cones probably takes place further down-stream in the degenerative pathway. Increasing lines of evidence suggest PKG as one of the main initiators of a photoreceptor cell death cascade ^4^, ^5^, together with increased Ca^2+^ levels ^39^. Excessive PKG activity could lead to the phosphorylation of a number of substrates associated with neuronal cell death, such as *Rac1* ^40^ or cyclic-AMP-response-element binding protein (CREB) ^41^. In addition, PKG phosphorylation dictates the activity of HDACs ^42^, in line with our previous observations that increased PKG and HDAC activities are temporally connected in 10 different animal models for inherited retinal degeneration ^4^. Since HDAC inhibition did not seem to affect VASP phosphorylation in cones, our study indicates that during cell death HDAC activity may occur independently of-or downstream of PKG. The underlying mechanisms through which HDAC inhibition offers cone protection remains to be determined.

While we found cone protection after late postnatal (PN14) HDAC inhibition, HDAC activity is also crucial for mouse photoreceptor development as HDAC inhibition at early postnatal retinal development (PN2) leads to a complete loss of developing rod photoreceptors ^43^. In mouse retina cone photoreceptors are born prenatally ^44^, even though the full maturation and commitment towards blue cones (S cones) or red/green cones (M cones) takes place during the second postnatal week ^45^. Since we performed HDAC inhibition at roughly the same time, we assessed the expression and localization of S- and M-opsin to address whether the observed increase in cone survival following HDAC inhibition may be due to cone development and maturation delays. In addition, we checked for the cone specific transducin (GNAT2) as a member of the cone phototransduction cascade ^46^. The observed correct localization of cone opsins and transducin suggest that late postnatal HDAC inhibition did not delay cone development and maturation. However, we cannot exclude the possibility that the rapid degeneration of *cpfl1* cones was in part caused also by developmental effects. In this case TSA treatment, at a critical period, may have enabled normal development as the observed effects were relatively long lasting (up to 10 days). Future studies may reveal whether TSA and HDAC inhibition mediated cone protection can be applied universally to cone degeneration in general, irrespective of developmental stage and disease pathogenesis.

In summary, we have shown for the first time that HDAC inhibition can prevent hereditary cone loss in *cpfl1* retina. At the same time HDAC inhibition improved cone migration without affecting cone differentiation and development. Our study provides a proof-of-principle highlighting HDAC inhibition as a relevant strategy for cone photoreceptor protection not only in hereditary cone dystrophies but also for other complex and more common cone diseases, such as diabetic retinopathy and age-related macular degeneration, associated with cone degeneration and improper cone migration.

## Materials and Methods

### Animals

*Cpfl1* and congenic C57BL/6 wild-type (wt) animals were housed under standard white cyclic lighting, had free access to food and water, and were used irrespective of gender. All procedures were approved by the Tübingen University committee on animal protection (Einrichtung für Tierschutz, Tierärztlichen Dienst und Labortierkunde) and performed in accordance with the ARVO statement for the use of animals in ophthalmic and visual research. Procedures performed at the polistab of University of Modena and Reggio Emilia (*in vivo* treatment on *cpfl1* and wt animals) were reviewed and approved by the local ethical committee (Prot. N. 50 12/03/2010). All efforts were made to minimize the number of animals used and their suffering. Because in *cpfl1* retina cone degeneration starts at postnatal day (PN) 14 and peaks at PN24 ^3^ all the analyses were performed at these ages.

### Immunofluorescence

Retinae were fixed in 4% paraformaldehyde (PFA) in 0.1 M phosphate buffer (pH 7.4) for 45 min at 4°C. Sagittal 12 μm sections were collected, air-dried and stored at −20°C. Sections were incubated overnight at 4°C with primary antibodies. The primary antibodies used were specific for Glycogen phosphorylase (1:1000 dilution, kindly provided by Prof. Pfeiffer-Guglielmi ^47^), M-opsin (1:200 dilution, AB5405, Millipore, Darmstadt, Germany), S-opsin (1:200 dilution, AB5407, Millipore), GNAT2 (1:200 dilution, sc-390, Santa Cruz, Dallas, USA), cGMP (1:500 dilution, kindly provided by Prof. Steinbusch, University of Maastricht, The Netherlands ^48^) and Phospho-VASP (Ser239) (1:200 dilution, 0047-100/VASP-16C2, Nanotools, Teningen, Germany ^3^, ^38^). Alexa Fluor 488- or 566-conjugated were used as secondary antibodies (Molecular Probes, Inc. Eugene, USA). Negative controls were carried out by omitting the primary antibody. Specificity of cone labeling with glycogen phosphorylase (Glyphos) was shown by co-labelling with Glyphos and peanut-agglutinin (see Supplementary Fig. S5), a well-established marker for cones ^49^.

### HDAC *in situ* activity assay

HDAC activity assays were performed on cryosections of 4% PFA- fixed eyes. The assay is based on an adaptation of the Fluor de Lys Fluorescent Assay System (Biomol, Hamburg, Germany). Retina sections were exposed to 200 μM Fluor de Lys-SIRT2 deacetylase substrate (Biomol) with 500 μM NAD+ (Biomol) in assay buffer (50 mM Tris/HCl, pH 8.0; 137 mM NaCl; 2.7 mM KCl; 1 mM MgCl2) for 2 hours at room temperature. Sections were then washed in PBS and fixed in methanol at −20°C for 20 min. x 0.5 developer (Biomol) in assay buffer was applied with 2 μM TSA (Sigma, Steinheim, Germany), 2 mM NAM (Sigma), and 500 μM NAD+ (Biomol) in assay buffer (50 mM Tris/HCl, pH 8.0; 137 mM NaCl; 2.7 mM KCl; 1 mM MgCl2)^7^. Negative controls consisted of omitting the substrate (Supplementary Figure 1e).

### Retinal explant cultures

Organotypic retinal cultures from *cpfl1* (n=7) and wt (n=4) animals that included the retinal pigment epithelium (RPE) were prepared under sterile conditions. Briefly, PN14 animals were sacrificed, the eyes enucleated and pretreated with 12% proteinase K (ICN Biomedicals Inc., OH, USA) for 15 minutes at 37°C in R16 serum free culture medium (Invitrogen Life Technologies, Paisley, UK). Proteinase K was blocked by addition of 10% fetal bovine serum, followed by rinsing in serum-free medium. Following this, cornea, lens, sclera and choroid were removed carefully, with only the RPE remaining attached to the retina. The explant was then cut into four wedges to give a clover-leaf like structure which was transferred to a culture membrane insert (Corning Life Sciences, Lowell, USA) with the RPE facing the membrane. The membrane inserts were placed into six well culture plates with R16 medium and incubated at 37°C in a humidified 5% CO_2_ incubator. The culture medium was changed every 2 days during the 10 culturing days.

Retinal explants were left without treatment for 2 days (until PN16), followed by Trichostatin A treatment (≥98% (HPLC), from Streptomyces sp., Sigma-Aldrich, St-Louis, USA) in two concentrations: 10 or 100 nM. Trichostatin A (TSA) was dissolved in 0.2% dimethyl sulfoxide (DMSO; Sigma) and diluted in R16 culture medium. For controls, the same amount of DMSO was diluted in culture medium. Culturing was stopped at PN24 by 2 h fixation in 4% PFA, cryoprotected with 30% sucrose and then embedded in tissue freezing medium (Leica Microsystems Nussloch GmbH, Nussloch, Germany).

### *In vivo* injections

Animals were anesthetized with an intraperitoneal injection of Ketamine (100ng/kg) Xylazin (5mg/kg) (Bela-Pharm, Vechta, Germany/Bayer Vital, Leverkusen, Germany), eye lids were anesthetized locally with Novesin (Omnivision, Puchheim, Germany) and animals were kept warm during injections. Single intravitreal injections were performed at PN14 on one eye while the other eye was sham injected with 0.0001% DMSO as contralateral control. TSA (1 and 10 nM) was diluted in 0.9% NaCl solution, which was also used for sham treatment. Injections were performed with 0.5 μl of 10 nM and 100 nm TSA in order to have a final concentration of 1 nM and 10 nM, respectively, assuming the free intraocular volume of mouse eye to be 5μl (http://prometheus.med.utah.edu/~marclab/protocols.html). Eleven *cpfl1* animals and eight wt from three different litters were used for intravitreal injections and were sacrificed 10 days after treatment (PN24). Eyes were immediately enucleated, fixed for 2 h in 4% PFA and prepared for cryosectioning (see ‘Retinal explant cultures’). Blinding was obtained by the analysis of the sections marked in a non-conclusive fashion.

### Microscopy, cell counting, and statistical analysis

Fluorescence microscopy was performed on an Axio Imager Z1 ApoTome Microscope, equipped with a Zeiss Axiocam digital camera. Images were captured using Zeiss Axiovision 4.7 software. Adobe Photoshop CS3 (Adobe Systems Incorporated, San Jose, CA) was used for primary image processing. To account for a difference in intravitreal injection sites, quantifications were performed on pictures captured on at least nine different random positions of at least three sagittal sections for at least three different animals for each genotype and treatment using Z-stacks mode of Axiovision 4.7 at 20x magnification. The average area occupied by a photoreceptor cell (*i.e*. cell size) for each individual eye was determined by counting DAPI-stained nuclei in 9 different areas (50 x 50 μm) of the retina. The total number of photoreceptor cells was estimated by dividing the outer nuclear layer (ONL) area by this average cell size. The quantification of cones was performed by manually counting the number of positively labelled cones in the ONL. Values obtained are given as fraction of total cell number in ONL (*i.e*. as percentage) and expressed as average ± standard error of the mean (SEM). For statistical comparisons the two-tailed, unpaired Student t-test as implemented in Prism 6 for Windows (GraphPad Software, La Jolla, CA) was employed.

To assess differences in cone migration, the distance in μm between the outer plexiform layer (OPL) and the center of Glyphos positive cell bodies was measured using Axiovision software (Zeiss). Distance values for 150-250 cones in the entire retinas were averaged on Glyphos immunostained sections from at least 5 different animals for each genotype. The migration profile of cones was presented as the relative migration distance to ONL thickness measured using Axiovision software. Statistical differences between experimental groups were calculated using Student’s two-tailed, unpaired t-test and Microsoft Excel software. Error bars in the figures indicate SEM, levels of significance were: * = *p<0.05*, ** = *p<0.01*, *** = *p<0.001*.

### Assessment of intravitreal clearance of TSA

The intravitreal clearance (CL_ivt_) of trichostatin A was calculated *in silico* using the Quantitative structure-property relationships (QSPR) model published recently ^50^ based on comprehensive rabbit data from intravitreal injection experiments: LogCL_ivt_= −0.25269 - 0.53747 (LogHD) + 0.05189 (LogD_7.4_), where HD is the number of hydrogen bond donor atoms and LogD_7.4_ is the calculated n-octanol/water distribution coefficient at pH 7.4 of the compound. The model is built based on small molecular weight compounds using the linear multivariate analysis tools: principal component analysis (PCA) and linear partial least square (PLS) (Simca Plus, version 10.5, Umetrics AB, Umea, Sweden). Firstly, the chemical structure of the TSA was retrieved from ACD/Dictionary from ACDlabs software (version 12, Advanced Chemistry Development, Inc., Toronto, Canada) and was used as input in ACDlabs software to generate 30 molecular descriptors: pK_a_ for the most acidic molecular form, pK_a_ for the most basic form, LogD at pH 5.5 and 7.4, LogP, MW, PSA (polar surface area), FRB (freely rotatable bonds), HD (hydrogen bond donors), HA (hydrogen bond acceptors), Htot (HD + HA), rule of 5, molar refractivity, molar volume, parachor, index of refraction, surface tension, density, polarizability, C ratio, N ratio, NO ratio, hetero ratio, halogen ratio, number of rings and number of aromatic, 3-, 4-, 5- and 6-membered rings. The applicability domain of the intravitreal clearance model for TSA was inspected generating the PCA score plot of the training set of the model together with TSA (Fig. S2). Compounds that lie inside the ellipse depicted in the plot belong to the same chemical space of the model and are predictable by the model. Once the applicability domain was confirmed, the intravitreal clearance value of TSA was calculated with the above equation using the descriptors values (LogD at pH 7.4 and HD). Half-life was calculated using equation t_1/2_ = ln2 V_d_/CL, where V_d_ is the volume of distribution and CL is the intravitreal clearance. V_d_ values of 1.18 – 2.18 ml were used, since this range covers 80% of the compounds ^50^.

## Acknowledgements

We are grateful to Prof. E. Zrenner for fruitful discussions and support, we also thank K. Masarini, and N. Rieger for skillful technical assistance and M. Power for critically reading of the manuscript. This work was supported by the Kerstan Foundation, Deutsche Forschungsgemeinschaft [DFG PA1751/4-1, DFG TR 1238 4-1], Alcon Research Institute, European Commission [DRUGSFORD: HEALTH-F2-2012-304963], German Ministry of Education and Research [BMBF HOPE2 – FKZ 01GM1108A]. We acknowledge support by Open Access Publishing Fund of University of Tübingen.

## Author contributions

D.T. and F.P-D conceived the study; D.T. and B.A-G. conducted the in vitro studies; D.T. and A.C. performed the in vivo studies; D.T. and M.B. conducted analysis of the in vitro treatment; D.T., M.B., M.K. and A.S. conducted analysis of the in vivo treatment; E.del A. and A.U. calculated in vivo clearance time; D.T., Y.A., V.M. and F.P-D. designed experiments and analyzed the data; S.M.H. and M.U. participated in results interpretation; D.T., Y.A., V.M. and F.P-D. wrote the paper. All authors have reviewed and approved the manuscript.

## Competing financial interests

The authors declare no competing financial interests.

